# Measuring Temporal Variations of Nucleotide Pools in Microbial Granular Biofilm Performing Enhanced Biological Phosphorous Removal

**DOI:** 10.1101/2025.08.27.672571

**Authors:** Jitske van Ede, Timothy Páez-Watson, Mark C.M. van Loosdrecht, Martin Pabst

**Author notes:** Corresponding authors: Timothy Páez-Watson, Jitske van Ede, Martin Pabst. These authors contributed equally to this work (Jitske van Ede, Timothy Páez-Watson).

## Abstract

Microbial communities often face environmental fluctuations that occur on timescales much shorter than their growth rate or proteome turnover. In such cases, cellular responses are likely driven by rapid changes in metabolite pools, particularly energy nucleotides including ATP, ADP, and AMP. However, robust methods to quantify these metabolites in biofilm-forming microbial communities are lacking. Here, we developed and systematically evaluated a metabolomics workflow for a granular biofilm enrichment, which performs Enhanced Biological Phosphorous Removal (EBPR). We combined fast quenching in liquid nitrogen and a boiling water extraction, followed by high resolution mass spectrometry, using porous graphitic carbon chromatography and ^13^C-labeled internal reference standards. Among tested procedures, a boiling water extraction was most suitable for extraction of nucleotides, as indicated by stable adenylate energy charge (AEC) and isotopic ratios. Applied to an anaerobic–aerobic cycle of a lab scale EBPR system, the method revealed dynamic changes in AEC and uridylate energy charge (UEC) during acetate uptake and polyphosphate degradation. These results demonstrate that energy pool imbalances underlie rapid metabolic switching observed in EBPR systems. Moreover, the established method provides a foundation for performing metabolomic studies of microbial biofilms in general.

## 1 Introduction

Microorganisms in natural and engineered environments continuously face fluctuations in physicochemical conditions, including changes in substrate availability^1^, pH^2^, temperature^3^, and redox potential^4^. To survive and adapt, cells employ regulatory responses that span multiple layers. The timescale of the environmental change relative to the cell’s response capabilities determines which layer of regulation dominates^5,6^. Transcriptomic responses and proteome remodelling typically require minutes to hours, especially considering that proteome remodelling is dependent on protein dilution by growth^7,8^. However, environmental perturbations can occur rapidly, and are often faster compared to proteome level changes^5,6^. Metabolic fluxes can also be regulated through allosteric regulation, post translational modifications and changes in substrate/product concentrations. Among these metabolites, nucleotides such as ATP, ADP, AMP, and their guanylate (GxP), uridylate (UxP), and cytidylate (CxP) counterparts are central. Their concentrations not only reflect the energy and redox status of the cell but also influence flux distributions and enzyme activity through thermodynamic and kinetic effects. Accordingly, metrics such as the adenylate energy charge (AEC)^10^ have long served as proxies for cellular energetic health, often maintained within a narrow physiological range of 0.7–0.95 across different species and conditions^11-14^.

An example where fast regulation clearly dominates is that of Enhanced Biological Phosphorous Removal (EBPR) processes^4^. The microbial community in this system experience rapid anaerobic–aerobic transitions during wastewater treatment cycles. Under anaerobic conditions, members of the community rapidly take up acetate and store it as polyhydroxyalkanoates (PHAs), fuelled by glycogen and polyphosphate degradation. Upon the introduction of oxygen, this metabolic strategy is abruptly reversed, with PHA oxidation supporting polyphosphate and glycogen replenishment^15^. These dynamic transitions occur within minutes, orders of magnitude faster than the imposed growth rates of the biomass (often referred to as sludge retention time) which span days. Consequently, proteome remodelling is too slow to explain the observed flux changes.

Recent proteomic studies revealed protein abundance changes of only < 3% during the EBPR cycle^24^, remarkably small considering the dramatic flux inversions observed. These observations have led to the hypothesis that core metabolic regulation underpinning EBPR transformations is driven by shifts in key metabolite pools within the cells of the community, particularly those associated with energy and redox metabolism^26,27^. Detailed metabolomic studies of EBPR communities have not been conducted to date, mainly due to two challenges: lack of pure cultures and the growth of the EBPR community as dense granular biofilms embedded in extracellular polymeric substances hampering metabolite extraction.

Various extraction methods have been explored for planktonic cells, including high-temperature extractions, freeze/thaw cycles, chloroform-methanol lysis and acid or alkaline treatments^28-33^. Biofilms often require an additional mechanical disruption such as bead beating and sonication to enhance metabolite recovery^34,35^. The choice of solvent also depends on the chemical characteristics of the target metabolites, including polarity and solubility^33^. As a result, there is no universal extraction method; protocols must be tailored to both the microbial system and metabolite class of interest. Moreover, metabolites like nucleotides turn over rapidly and require instant quenching to preserve their in vivo state^30,36^. Quenching strategies typically include a solvent solution combined with extreme temperatures or pH^30^, such as cold methanol^31,37^, liquid nitrogen^33,38^ or acidic organic solvents^28^.

Two key factors are commonly evaluated when testing the extraction efficiency: efficacy and stability^30^. The efficacy is assessed by applying different extraction methods to the same biological sample and comparing their ability to extract different metabolites. On the other hand, stability can be evaluated by spiking metabolite standards and measuring recovery rates^30^. While metabolite degradation should generally be prevented, minor changes can be corrected for by using internal standards^31^. Isotope dilution mass spectrometry uses a uniformly ^13^C-labeled cell extract as an internal standard, which allows to correct for degradation during sample preparation, injection variability, ion suppression effects in electrospray ionisation, and instrumental fluctuations^33,39^.

Metabolomics is predominantly analysed via liquid chromatography coupled to mass spectrometry via electrospray ionisation (ESI), due to its high sensitivity, selectivity and dynamic range compared to enzymatic assays or UV and fluorescence detection^40^. Moreover, it allows for untargeted analysis of metabolites and requires only minute amounts of sample. Earlier techniques to separate nucleotides include ion pair reverse phase (IPRP) and ion exchange chromatography (IEC)^41^ . While the latter is only poorly compatible with ESI-MS due to high salt concentrations, IPRP is not ideal due to persistent contamination by IP reagents^42-44^. On the other hand, hydrophilic interaction liquid chromatography (HILIC) is compatible with ESI-MS and effective for separating nucleotides. However, it requires a high proportion of organic solvent, which can hinder the solubility of some metabolite extracts. HILIC also often suffers from peak broadening and retention time drift, affecting method robustness and reproducibility, though certain columns have been shown to diminish these issues^4,24,25^. A promising alternative is porous graphitic carbon (PGC) chromatography, which shows great separation performance for polar metabolites and is compatible with ESI-MS^44-47^. Nevertheless, PGC can suffer from retention time instability and column polarization, which, however, can be largely circumvented by using internal standards, thorough sample cleanup, choice of running buffers and additives, and electrical grounding^45,48,49^.

Here, we demonstrate a robust method for quantifying nucleotide pools in a granular biofilm microbial community performing EBPR. By combining rapid quenching with liquid nitrogen, boiling water extraction, PGC chromatographic separation followed by high resolution mass spectrometric detection, we were able to capture energy nucleotide dynamics during a typical EBPR anaerobic–aerobic cycle. Our findings provide evidence for metabolite-level regulation and highlight the potential of energy charge modulation in driving the EBPR transformations.

## 2 Materials and Methods

### EBPR granular sludge enrichment

A lab scale EBPR system was run in a 1L (0,8 L working volume) sequencing batch reactor (SBR), following conditions described in Páez-Watson, et al. ^50^ with some adaptations. The reactor was inoculated using enriched sludge from the work of Páez-Watson, et al. ^50^, which was previously inoculated with activated sludge from a municipal wastewater treatment plant (Harnaschpolder, The Netherlands). Each SBR cycle lasted 6 hours, consisting of 10 minutes of settling, 70 minutes of effluent removal, 10 minutes of N_2_ sparging, 5 minutes of anaerobic feeding, 130 minutes of anaerobic phase and 135 minutes of aerobic phase. N_2_ gas and compressed air were sparged at 500 ml/min into the reactor broth to maintain anaerobic and aerobic conditions respectively. The hydraulic retention time (HRT) was 12 hours (removal of 400 ml of broth per cycle, each cycle of 6 hours). The average solids retention time (SRT) was controlled to 8 days by the removal of 25 ml of mixed broth at the end of the mixed aerobic phase in each cycle (total 100 ml per day). The pH was controlled at 7.3 ± 0.1 by dosing 0.5 M HCl or 0.5 M NaOH. The temperature was maintained at 20 ± 1 °C.

The reactor was fed with three separate media components diluted in demineralized water: a concentrated COD medium (400 mg COD/L) of acetate (17 g/L sodium acetate ×3H_2_O); a concentrated mineral medium (2.51 g/L NaH_2_PO4×H_2_O, 1.53 g/L NH_4_Cl, 1.59 g/L MgSO_4_×7H_2_O, 0.4 g/L CaCl_2_×2H_2_O, 0.48 g/L KCl, 0.4 g/L N-allylthiourea (ATU), and 6 mL/L of trace element solution prepared following Smolders, et al. ^15^); and demineralized water containing 5 mg/L of yeast extract. In each cycle, 40 mL of COD medium, 40 mL of mineral medium and 320 mL of water with yeast extract were added to the reactor during the 5 minutes of feeding. The final feed contained 400 mg COD/L of acetate.

### Reactor and biomass analyses

Extracellular concentrations of phosphate and ammonium were measured with a Gallery Discrete Analyzer (Thermo Fisher Scientific, Waltham, MA). Acetate was measured by high performance liquid chromatography (HPLC) with an Aminex HPX-87H column (Bio-Rad, Hercules, CA), coupled to RI and UV detectors (Waters, Milford, MA), using 0.0015 M phosphoric acid as eluent supplied at a flowrate of 1 mL/min.

The biomass concentration (total and volatile suspended solids – TSS and VSS) was measured in accordance with Standard Methods as described in Smolders, et al. ^15^ with some modifications: 10 mL of mixed broth were obtained at the end of the aerobic phase, centrifuged at 3600*g for 3 minutes and washed twice with demineralized water to remove salts. The sludge was then dried at 100 °C for 24 hours and weighed on a microbalance to determine the dry content – TSS. The ash content was determined by incinerating the dry material in an oven at 550 °C, and the difference used to calculate the VSS.

For glycogen and PHA determination, biomass samples (10 mL mixed broth) were collected throughout the batch test and stored in 15 ml conical tubes containing 0.3 mL of 37 % formaldehyde to stop biological activity. After each batch test, the biomass tubes were pottered to break the granular structure of the biomass, centrifuged at 3700 *g for 5 minutes and washed twice. The pellet was then frozen at -80°C for at least 3 hours and freeze dried. For glycogen analysis the method described by Smolders, et al. ^15^ was used: 5 mg of dry biomass was digested in 0.9 M HCl solutions in glass tubes at 100 °C for 5 hours. After this time, tubes were cooled at room temperature, filtered with 0.45 µm Whatman disk filters and neutralized with equal volumes of 0.9 M NaOH. The glucose resulting from digestion was quantified using the D-Glucose Assay Kit (GOPOD Format) from Megazyme (Bray, Ireland).

For PHA extraction the method described by Oehmen, et al. ^51^ was used. Approximately 30 mg of freeze-dried biomass was placed in a tube with 1.5 mL of 10% sulfuric acid in methanol and 1.5 mL of chloroform. To each tube, 100 μL of an internal standard solution (0.01 g/mL benzoic acid in methanol) was added. The samples were then hydrolyzed and esterified at 100 °C for 5 hours with periodic manual vortexing. Following this, 3 mL of ultrapure water was added to facilitate phase separation. The organic phase containing the esters was filtered and analyzed using a gas chromatograph (GC 6890 N, Agilent, USA). Quantifications of PHB, PH2MV, and PHV were performed using commercial standards, including 3-hydroxybutyrate, 2-hydroxyhexanoate, and a synthetic copolymer of (R)-3-hydroxybutyrate-(R)-3-hydroxyvalerate (Sigma-Aldrich, USA).

### Whole metagenome sequencing

For metagenomics, DNA from the biomass samples was extracted using the DNeasy PowerSoil Pro-Kit (Qiagen, Germany) following the manufacturer’s protocol. Shotgun sequencing was performed by Hologenomix (Delft, The Netherlands). Paired-end sequencing with a read length of 150 bp was conducted using the Illumina NovaSeq X sequencing system. Library preparation was carried out using the Nextera XT DNA Library Preparation Kit. Approximately 10 Gbp of sequencing data were generated per sample. The quality of raw sequenced reads was assessed using FastQC (version 0.11.7) with default parameters^52^, and results were visualized with MultiQC (version 1.19). Low-quality paired-end reads were trimmed and filtered using Fastp (version 0.23.4) in paired-end mode^53^.

Taxonomic profiling from raw reads was performed by Hologenomix (Delft, The Netherlands) using the standard Kraken2 (version 2) database (using all complete bacterial, archaeal, and viral genomes in NCBI Refseq database, complemented with a curated wastewater database) with default parameters^54^. The taxonomic classification outcomes from Kraken2.0 were converted into abundance tables using the biom file converter tool to explore metagenomics classification datasets. Figures were generated with the R package “phyloseq”^55^. Clean reads were further assembled into contigs using MetaSPAdes (version 3.15.5) with default parameters^56^. From there, protein-coding sequences were predicted from contigs using Prodigal (https://github.com/hyattpd/Prodigal). The resulting fasta file was further used for metaproteomic analysis. Sequencing data are available through the BioProject database https://www.ncbi.nlm.nih.gov/bioproject/ under the BioProject no. PRJNA1305385.

### Metaproteomics

Samples were taken during the aerobic phase of the anaerobic-aerobic cycle as described in the section “Sampling and quenching for Metabolomics”. The crushed sample was transferred to a LoBind Eppendorf tube, freeze-dried overnight and subsequently processed as described in van Olst et al. (2025)^57^. In short, approximately 3.5 mg of biomass (dry weight) was lysed by three rounds of vortexing at max speed with acid washed glass beads after the addition of B-PER reagent and 50 mM TEAB pH 8. The biomass was heated to 80 °C in the Thermomixer with Thermotop (1000 rpm), and sonicated for 10 min in an ultrasonic bath. After pelleting the biomass, the supernatant was collected in a clean LoBind Eppendorf tube and proteins were precipitated by the addition of TCA (ratio 4:1) at 4 ° for 30 min. The sample was centrifuged at 14000 rcf, 4 °C for 15 min and the protein pellet was washed with ice cold acetone, centrifuged for 3 min, 14000 rcf, and dissolved in 100 μL 6 M urea. Reduction was performed by the addition of 30 μL 10 mM dithiothreitol at 37 °C for 60 min in the Thermomixer with Thermotop (300 rpm), and alkylation was performed by adding 30 μL 20 mM iodoacetamide at room temperature in the dark for 30 min. Subsequently, 100 mM ammonium bicarbonate was added to dilute the Urea concentration to <1 M and the sample was digested with 15 μL 0.1 μg/μL Trypsin for approximately 18 hours at 37 °C, 300 rpm. A solid phase extraction was performed as described by van Olst et al. (2025)^57^. Finally, the sample was speed vac dried and dissolved in 3% acetonitrile, 0.1% formic acid in water to reach a concentration of 0.5 mg/mL, where approximately 500 ng was analysed by shotgun proteomics.

Shotgun proteomics was performed on an EASY 1200 nano-LC connected to a Q Exactive plus Orbitrap Mass Spectrometer (Thermo Scientific, Germany). Peptides were separated on a C18 PepMap column (0.05 x 150 mm column, Thermo Scientific) using a constant flow rate of 350 nL/min, with mobile phase A (0.1% formic acid (FA), 1% acetonitrile (ACN) in Optima LC/MS-grade H2O) and mobile phase B (0.1% FA, 80% ACN in Optima LC/MS-grade H2O). The LC gradient included an initial 2 min at 5% B, followed by a linear increase to 25% B over 90 min, a linear increase to 55% B over 60 min, then a return to 5% B within 3 min, and an equilibration for an additional 20 min. Electrospray ionization was carried out in positive mode. MS1 spectra were acquired at a resolution of 70,000, an automatic gain control (AGC) of 3e6 and a maximum injection time of 75 ms. Data-dependent acquisition (DDA) was applied, isolating precursors using a 2.0 m/z isolation window over a scan range of 385-1250 m/z. Fragmentation was performed with a normalized collision energy of 28%. MS2 spectra were acquired at a resolution of 17,500, an AGC of 2e5 and an maximum injection time of 75 ms. Database searching was performed in PEAKS Studio 10.5 (Bioinformatics Solutions Inc.) using a reference sequence database obtained from whole metagenome sequencing, with a parent mass error tolerance set to 20.0 ppm, a fragment mass error tolerance set to 0.02 Da and Trypsin as the proteolytic enzyme. Carbamidomethylation was specified as a fixed modification, while oxidation and deamidation were set as variable modifications, allowing up to three variable modifications per peptide. The identified peptide spectrum match list was furthermore taxonomically annotated using the NovoBridge pipeline^58^, which retrieves a taxonomic lineage from the Unipept database (https://unipept.ugent.be/) for every peptide sequence, and which are then grouped into a taxonomic composition for every taxonomic rank. Raw shotgun metaproteomics data, the reference sequence database and the search files are available in the PRIDE database under project accession number PXD067586.

### Sampling and quenching for Metabolomics

Samples were collected from the stirred broth of the bioreactor using a 5 mL syringe attached to the sampling port. For method optimization, samples were taken at the end of the aerobic phase, whereas for dynamic profiling, samples were collected throughout the anaerobic–aerobic cycle. Each sample, containing both broth and biomass granules, was immediately transferred into a pre-chilled mortar containing liquid nitrogen, ensuring submersion within 3 seconds of collection. A precooled pestle was then used to crush the frozen sample into a fine powder in the presence of excess liquid nitrogen. The resulting powder was transferred into 2 mL Eppendorf tubes with manually pre-cut lids to allow them to fit inside 15 mL tubes used during heat-based extraction methods, and 50 µL of uniformly ^13^C-labeled yeast extract was added as an internal standard. The ^13^C-labeled yeast extract was prepared in house as published by Wu Liang et al., 2005^39^. The tubes were stored in liquid nitrogen until further processing. Depending on the quenching and extraction protocol, one of the following procedures was used to inactivate metabolic enzymes and prepare the samples: <u>Acidic Acetonitrile/Methanol</u>: 500 µL of pre-cooled (at –20 °C) acetonitrile:methanol:water (40:40:20, v/v/v) with 0.1 M formic acid was added to the sample. After vortexing, samples were placed on ice for 3 minutes. Subsequently, 43.5 µL of pre-cooled 15% (w/v) ammonium bicarbonate in MS-grade water was added to neutralize the extract. Samples were then stored at −80 °C until further processing by solid-phase extraction (SPE) and LC-MS analysis.

<u>Boiling Ethanol</u>: 2 mL of molecular-grade ethanol was preheated in 15 mL conical tubes using a 100 °C water bath. One by one, frozen samples were removed from liquid nitrogen, lids were discarded, and tubes were immediately submerged in boiling ethanol. After brief mixing by inversion, samples were incubated in the water bath for 3 minutes, then cooled on ice and transferred to fresh 2 mL Eppendorf tubes for further processing.

<u>Boiling Water</u>: 2 mL of miliQ water was added to 15 mL conical tubes and preheated in a 100 °C water bath. Frozen samples were retrieved individually from liquid nitrogen, the pre-cut lids were removed, and the tubes were immediately submerged in the boiling water. After brief mixing by inversion, the samples were incubated at 100 °C for 3 minutes to ensure enzyme inactivation. Tubes were then cooled on ice and transferred to fresh 2 mL Eppendorf tubes for subsequent processing.

### Solid phase extraction of metabolites

Sample were centrifuged for 15 minutes at 14000 rcf and 4 °C using the Eppendorf centrifuge 5424R, after which the supernatant was transferred to a clean Eppendorf tube. The samples were subsequently washed and concentrated with a HyperSep^TM^ Hypercarb^TM^ SPE 96-well plate (25/1 ml Plate, Thermo Scientific). In short, the columns were preconditioned by washing with 600 μL 10 mM ammonium bicarbonate (ABC) in 60% acetonitrile (ACN), followed by an equilibration with 2 x 600 μL MS-grade H_2_O. The samples were loaded onto the columns and a wash step with 1 ml MS-grade H_2_O was performed. The samples were eluted using 2 x 200 μL 10 mM ABC in 60 % ACN. The samples were collected in clean LoBind Eppendorf tubes and dried using the SpeedVac concentrator plus (Eppendorf) at room temperature. The dried samples were redissolved in 20 μL MS-grade H_2_O prior to analysis by LC-MS.

### PGC-HRMS metabolomics

Liquid chromatographic separation coupled to high resolution mass spectrometric detection was conducted using an Acquity Ultra Performance Liquid Chromatograph (Waters, UK) coupled to a Q Exactive™ Focus Hybrid Quadrupole-Orbitrap™ Mass Spectrometer (Thermo Scientific, Germany). Chromatographic separation was achieved on a 100 x 1 mm Hypercarb™ Porous Graphitic Carbon HPLC Column (Thermo Scientific, No. 35005-101030) using a constant flow rate of 100 µL/min. The mobile phases consisted of 50 mM ammonium bicarbonate in MS-grade H_2_O (Phase A) and 100% MS-grade acetonitrile (Phase B). A linear gradient from 2.5% to 35% Phase B was applied over 18 minutes, followed by a gradient increase to 60% B over the next 6 minutes. The system was then returned to the initial conditions and equilibrated for 4 minutes. Each sample was analysed in duplicate using 6 μL injections, followed by two blank runs. High resolution mass spectrometric detection was performed in negative ion mode, scanning across a mass range of 100–850 m/z. The mass spectrometer was set to a resolution of 35K, with an AGC target of 1.0E6 and a maximum injection time of 100 ms. Raw data were processed using XCalibur 4.1 (Thermo Fisher Scientific, Germany). In addition, raw MS files were converted to mzXML files using MSConvertGUI, and the integration of nucleotides was performed in Matlab R2022a (MathWorks). Calibration of the mass spectrometer was performed using the Pierce™ LTQ ESI negative ion calibration solution (Thermo Fisher Scientific, Germany).

## 3 Results

### 3.1 Establishing a metabolomics pipeline for nucleotide quantification in EBPR granular biomass

A stable isotope dilution mass spectrometry pipeline was developed for quantifying the nucleotides AMP, ADP, ATP, UMP, UDP, UTP, GMP, GDP, GTP, CMP, CDP and CTP in a granular biofilm from a lab scale EBPR system. This included homogeneous biomass sampling, selecting an effective quenching and metabolite extraction method, establishing an effective chromatographic separation of nucleotides and validating possible interference from extracellular nucleotides (Figure 1A). Finally, we demonstrated the method by monitoring nucleotide concentrations throughout an EBPR anaerobic-aerobic cycle.

**Figure 1.**
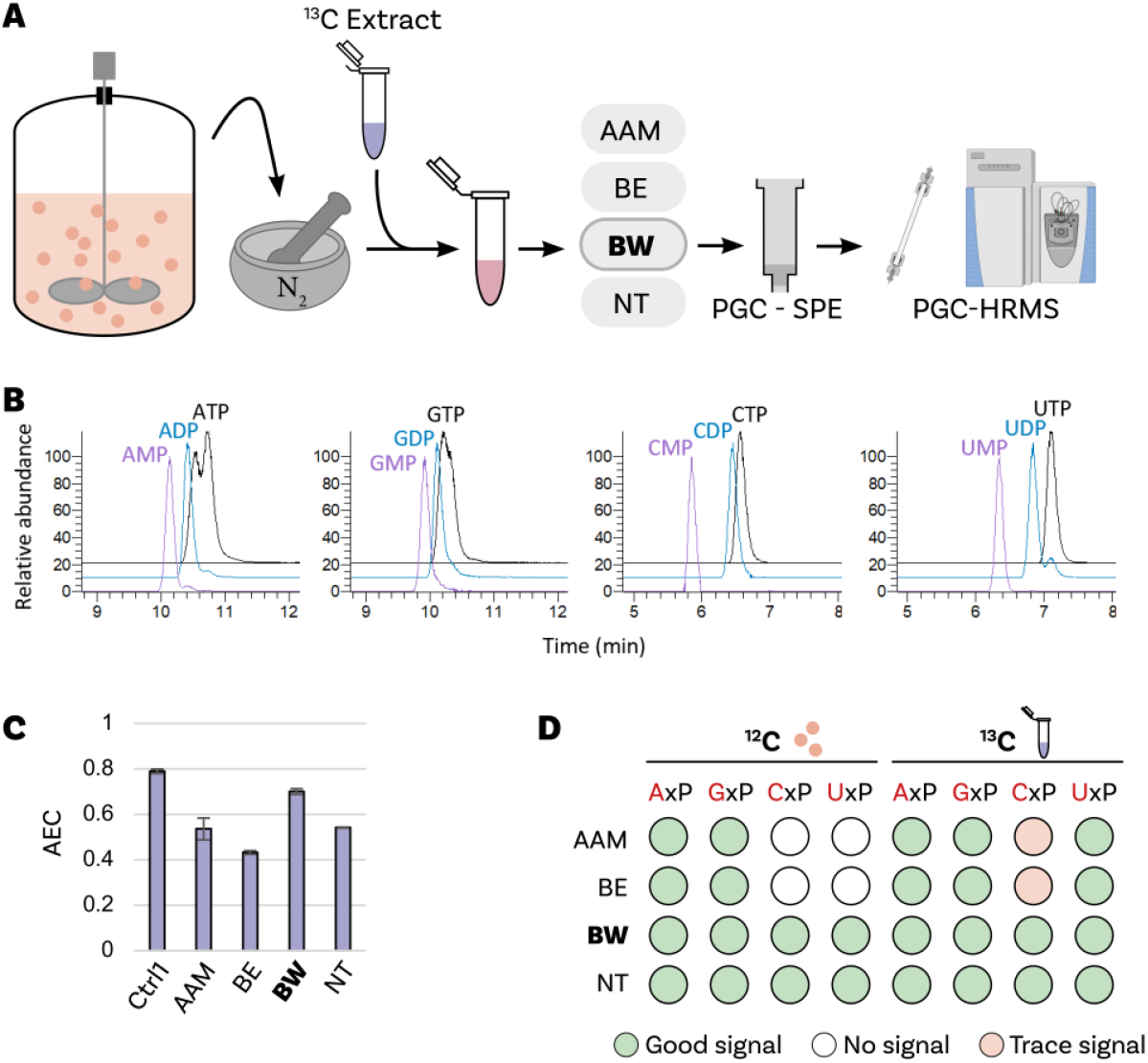
**A)** Schematic overview of the workflow to quantify nucleotides in a granular biofilm microbial community performing EBPR. Broth samples of the lab scale EBPR system were frozen in liquid nitrogen and crushed with a mortar and pestle. The collected samples were spiked with uniformly ^13^C-labeled yeast extract followed by either an acidic acetonitrile methanol extraction (**AAM**), a boiling ethanol extraction (**BE**), a boiling water extraction (**BW**) or no treatment (**NT**). A solid phase extraction (SPE) was performed followed by a chromatographic separation on porous graphite carbon (PGC) and high resolution mass spectrometric analysis. **B)** Extracted ion chromatograms of the uniformly ^13^C-labelled nucleotides from the analysed ^13^C yeast extract. **C)** Adenylate energy charge (AEC) calculated from the ^13^C nucleotide PGC-HRMS signals of the spiked ^13^C yeast extract, after performing different extraction approaches. AEC = ([ATP] + 0.5[ADP])/([ATP] + [ADP] + [AMP]). **D)** Detectability of the ^12^C nucleotides and the ^13^C isotope variants after performing various extraction methods. **Ctrl1:** ^13^C yeast extract.

The separation prior to mass spectrometric detection was performed using porous graphitic carbon (PGC) as the chromatographic separation phase combined with ammonium bicarbonate as mobile phase. PGC was chosen for its previously demonstrated effectiveness in retaining and separating nucleotides using volatile buffers^45-47^. To address common issues such as retention time instability and column polarization^48,49^, the ammonium bicarbonate concentration in the mobile phase was increased to 50 mM, and electrical grounding points were installed before and after the separation column. As a result, interday retention time shifts were limited to ±1 minute. Moreover, the PGC column enabled the separation of nucleotides in the order of mono-, di-, and tri-phosphates, allowing to estimate the contribution of in-source fragmentation to the abundance of analysed metabolites (Figure 1B).

Calibration curves of all nucleotides were established using synthetic reference standards spiked with a ^13^C-labeled yeast extract^39^, to allow determination of absolute nucleotide concentrations. The ^12^C/^13^C signal intensities of each nucleotide was plotted against the known ^12^C nucleotide concentrations. All calibration curves demonstrated strong linearity (R^2^ approaching 1) within the concentration range of 0.7-2150 μM (Table 1, Figure S4).

**Table 1.** Signal response curves for quantified metabolites by plotting the ^12^C/^13^C signal intensity of each metabolite against its ^12^C metabolite concentration. Linear fits were forced through zero, covering a concentration range of 0.7–2150 μM, where the squared correlation coefficient (R^2^) is provided, confirming excellent linearity. In addition, intra- and extracellular metabolite concentrations of the granular biofilm microbial community are provided. The concentrations are averages of two biological duplicates, which were each analysed twice (technical duplicates). The standard deviation is provided (+/-standard deviation). For all nucleotides, the extracellular concentration was < 1%.

|  | **R<sup>2</sup> for**<br>**metabolite**<br>**range: 0.7–2150**<br>**$\mu\text{M}$** | **Broth nucleotides**<br>**($\mu\text{M}$)** | **Extracellular**<br>**nucleotides**<br>**($\mu\text{M}$)** | **Extracellular**<br>**nucleotides**<br>**(%)** | **Average nucleotide**<br>**concentration**<br>**(mmol/mg X)** |
| --- | --- | --- | --- | --- | --- |
| ATP | 0.9931 | 58.1 +/- 3.6 | 0.48 +/- 0.07 | 0.83 | 1.51 (+/- 0.30) |
| ADP | 0.9919 | 63.0 +/- 1.8 | 0.48 +/- 0.06 | 0.76 | 0.78 (+/- 0.11) |
| AMP | 0.9905 | 84.5 +/- 2.1 | 0.66 +/- 0.08 | 0.78 | 0.45 (+/- 0.13) |
| GTP | 0.9889 | 39.3 +/- 3.6 | 0.14 +/- 0.02 | 0.35 | 0.67 (+/- 0.12) |
| GDP | 0.9904 | 21.3 +/- 0.5 | 0.09 +/- 0.02 | 0.40 | 0.20 (+/- 0.02) |
| GMP | 0.9880 | 26.5 +/- 0.8 | 0.17 +/- 0.03 | 0.64 | 0.28 (+/- 0.06) |
| CTP | 0.9841 | 134.7 +/- 0.4 | 0.10 +/- 0.06 | 0.08 | 2.25 (+/- 0.54) |
| CDP | 0.8784 | 9.9 +/- 0.2 | 0.00 +/- 0.003 | 0.03 | 0.09 (+/- 0.02) |
| CMP | 0.9926 | 7.5 +/- 0.3 | 0.03 +/- 0.001 | 0.40 | 0.06 (+/-) 0.02 |
| UTP | 0.9932 | 64.2 +/- 0.2 | 0.34 +/- 0.04 | 0.53 | 1.44 (+/- 0.31) |
| UDP | 0.9658 | 37.0 +/- 0.9 | 0.17 +/- 0.003 | 0.47 | 0.37 (+/- 0.08) |
| UMP | 0.9804 | 24.9 +/- 2.0 | 0.00 +/- 0.00 | 0.00 | 0.17 (+/- 0.06) |

In order to assess the sampling procedure, stirred-tank bioreactor samples were immediately immersed in liquid nitrogen and crushed into a fine powder, both to quench the metabolism as well as to facilitate metabolite extraction. This approach was evaluated and shown to ensure homogeneous biomass sampling from the bioreactor, yielding an average biomass concentration of 6.77 ± 0.06 g/L (triplicate samples).

Subsequently, four metabolite extraction methods were evaluated, including an acidic acetonitrile-methanol extraction followed by neutralization (AAM), a boiling ethanol extraction (BE), a boiling water extraction (BW), and one without additional treatment (NT). To assess metabolite conversion and degradation, all samples were spiked with ^13^C-labeled yeast extract^39^. The adenylate energy charge 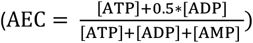 of the ^13^C-labeled metabolites in the samples following the different extraction procedures was compared to that of an untreated ^13^C-labeled yeast extract (control). Among the extraction methods, the BW method yielded an AEC of 0.70 ± 0.01, which was closest to the control value of 0.79 ± 0.01 (Figure 1C). In contrast, the AEC for samples extracted using AAM, BE, and NT showed a more pronounced decrease, with values of 0.54 ± 0.05, 0.43 ± 0.01, and 0.54 ± 0.002, respectively. In addition, all possible ^13^C nucleotide ratios (NTP/NDP, NDP/NMP, and NTP/NMP) were compared against the control sample. The NTP/NDP ratios for all nucleotides followed a similar pattern to that seen for the AEC, with the BW method producing values closest to the control (Figure S2). However, although the BW method also yielded NTP/NMP and NDP/NMP ratios that were generally closest to the control, these ratios were often significantly lower than those of the control sample (Figure S2). This indicates that still some conversion/degradation of the cellular metabolome was ongoing, likely increasing the levels of NMPs. Further, the ^12^C nucleotides were analysed for the different extraction procedures (Figure S3). Only trace signals were detected for CxP and UxP (x representing mono-, di-, or tri-phosphate) in samples extracted using the AAM and BE method (Figure 1D, Figure S3), while a good signal was obtained for the BW method. Since the BW extraction introduced the least bias in the^13^C AEC and ^13^C nucleotide ratios, while also providing ^12^C signals for all nucleotides of interest, this method was chosen for the remainder of the study.

Furthermore, to assess potential metabolite degradation during sample storage in the autosampler prior to analysis, ^13^C-labeled yeast extract was injected in regular time frames throughout the analysis period. Both the AEC and nucleotide ratios were monitored, revealing that the AEC and most nucleotide ratios remained stable over time, indicating minimal to no nucleotide degradation. This control was performed during all analyses (Figure S1.1, S1.2, and S1.3).

Finally, since samples are immediately frozen and pulverized upon collection, no distinction is made between intra- and extracellular nucleotides. This method was chosen to minimize sampling time and capture the metabolome as accurately as possible at a specific time point. Therefore, no separation of cell biomass from the supernatant was performed. To support this choice, an additional evaluation was conducted to assess whether extracellular nucleotide levels interfere with the quantification. Alongside the metabolite extraction of a regular broth sample (biomass in medium), an extraction was performed on a filtered broth sample, representing the extracellular environment. Extracellular nucleotide concentrations were found to contribute to less than 1% to the total broth concentration (Table 1), confirming that extracellular metabolite levels are negligible.

### 3.2. Variations in the nucleotide pool of PAOs in response to the environment

The established metabolomics method was employed to a typical anaerobic / aerobic EBPR cycle enriched with granular biomass. The microbial community contained various bacterial groups, with *Candidatus* Contendobacter and *Dechloromonas* showing the highest relative abundance based on taxonomic profiling using metaproteomics (Figure 2C). A characteristic EBPR profile was observed during the cycle^15^. In the initial anaerobic phase (first 30 minutes), acetate uptake coincided with the degradation of polyphosphate and glycogen, resulting in a P/C ratio of 0.376 P-mol released per C-mol uptake. Simultaneously, polyhydroxyalkaonate (PHA) accumulation occurred, predominantly as polyhydroxybutyrate (PHB), with a smaller fraction of polyhydroxyvalerate (PHV). During the aerobic phase, these PHAs were consumed, leading to the uptake of phosphate from the media and the replenishment of polyphosphate and glycogen (Figure 2B).

**Figure 2.**
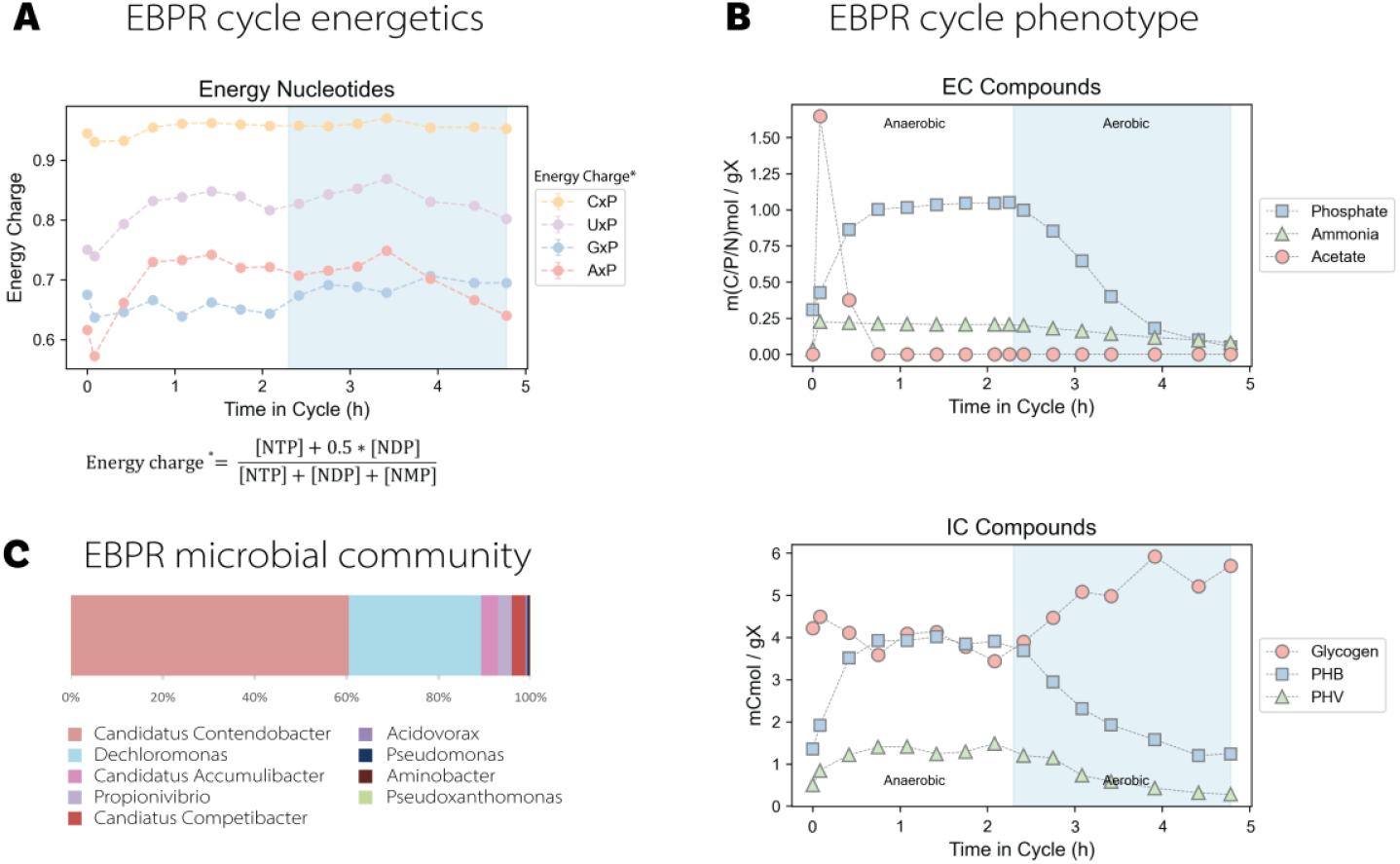
Anaerobic/aerobic cycle of a granular biofilm microbial community performing EBPR fed with acetate. **A)** The energy balance for AxP, GxP, CxP and UxP was calculated by: energy balance = ([NTP] + 0.5[NDP])/([NTP] + [NDP] + [NMP]). **B)** The phosphate, ammonia and acetate concentrations are presented in m(C/P/N)mol/gX, and the glycogen, PHB and PHV concentrations in mCmol/gX. **EC:** extracellular; **IC:** intracellular. **C)** The composition of the microbial community as determined by metaproteomics using NovoBridge^58^.

Quantification of nucleotide energy charges revealed distinct dynamics across the cycle (Figure 2A). The adenylate energy charge (AEC), calculated as ([ATP] + 0.5[ADP])/([ATP] + [ADP] + [AMP]), increased during the first hour of the anaerobic phase, reaching a peak of approximately 0.72. This increase coincided with acetate uptake and phosphate release. Following acetate depletion, the AEC plateaued for the remainder of the anaerobic phase. Upon transition to aerobic conditions, a minor transient increase was observed, followed by a gradual decline in AEC, returning to the initial value of ∼0.6 by the end of the cycle.

The uridylate energy charge (UEC), calculated analogously as ([UTP] + 0.5[UDP])/([UTP] + [UDP] + [UMP]), followed a similar trend to AEC, with a peak during the anaerobic phase and a gradual decline during the aerobic phase. The guanylate energy charge (GEC) showed modest variation, with a slight increase during the aerobic phase. In contrast, the cytidylate energy charge (CEC) remained consistently high and stable throughout the entire cycle, with minimal differences between the anaerobic and aerobic phases. Together, the data show that nucleotide energy charge varies over the EBPR cycle, with adenylate and uridylate pools exhibiting more pronounced temporal dynamics than guanylate and cytidylate pools.

## 4 Discussion

In this study, we established a targeted and quantitative workflow to quantify nucleotide pools in granular biofilms, as exemplified for a microbial community performing EBPR. In the following, we elaborate on the strengths and potential limitations of the established pipeline. Subsequently, we discuss the observed nucleotide dynamics in relation to EBPR transformations during anaerobic–aerobic cycling.

### 4.1 A quantitative nucleotide metabolomics pipeline for granular biofilms

Small changes in energy nucleotides can significantly impact metabolic flows, thereby holding valuable information. Although numerous metabolomics studies have been conducted on planktonic cells, limited method development has been dedicated to granules or biofilms. Most of these efforts predominantly focused on untargeted methods to maximize metabolite identification^59^. While useful for comparing planktonic and biofilm states or distinguishing bacterial species^34,35,60^, these approaches lack the precision required for quantitative metabolome analyses in granules and biofilms. Moreover, common pre-quenching sample handling steps, including centrifugation and washing, can distort the metabolite ratios and should be avoided^36,38^.

The presented method enables rapid, homogeneous biomass freezing. Due to the speed of metabolite conversions, minor environmental shifts may still alter the metabolome; such conversions could be reduced by even faster sampling. In yeast research, rapid sampling devices have reduced sampling times to <1 second^61^. Nevertheless, the tubes and sampling ports required for high speed are too small for working with larger aggregates of biomass such as granules.

Among the tested extraction methods, the BE extraction proved inadequate (Figure 1C), possibly due to a poor recovery of phosphorylated compounds, as previously reported^62^. Both the NT and AAM extractions resulted in altered nucleotide profiles compared to the control, indicating potential degradation or residual enzymatic activity. Although other studies have reported improved triphosphate recovery using similar cold acidic^38^ or acidic acetonitrile/methanol/water (40:40:20)^28^ extractions, those analyses were performed on planktonic *E. coli* cells and did also not include an extraction with boiling water. The structurally distinct biomass used in this study (granules) may account for differences in extraction efficiency and possible degradation processes.

The BW extraction method proved to be the most effective (Figure 1C), yielding ^13^C AEC and NTP/NDP ratios closest to the control. A hot water extraction has previously enabled the quantification of over 50 hydrophilic metabolites in planktonic *E. coli*^63^, including nucleotides, yet that study did include a centrifugation step, which may alter metabolite ratios and should be avoided. Despite the effectiveness of the BW method, deviations in the NDP/NMP and NTP/NMP ratios were observed compared to the control, suggesting the presence of residual enzymatic activity, metabolite degradation and/or matrix effects during MS analysis. Although the spiked ^13^C yeast extract can account for some of these factors, it cannot correct for enzymatic interconversion between nucleotides, unless the nucleotide concentrations in the spiked standard closely match those in the actual sample. This limitation could be addressed by generating a uniformly ^13^C-labeled extract from the enrichment culture of interest, thereby replacing the ^13^C yeast extract in the experiments. As such, the current method requires cautious interpretation of the NDP/NMP and NTP/NMP ratios.

Previous research has shown that metabolite profiles can differ between bacteria growing in planktonic form and those in a biofilm state^34,35,60^. An intriguing question that remains is whether bacteria also exhibit distinct metabolic profiles based on their spatial location, such as the surface versus the core of a granule. These differences could arise from variations in access to nutrients and oxygen, influenced by factors like granule size, density, and the diffusion coefficient of solutes^64^. Smaller granules are likely to have a more uniform metabolome. The current method averages measurements across the entire bacterial population, potentially masking spatial variations. Future advancements might help to address these type of questions. Nevertheless, the employed approach reveals how the energy state changes when varying the environmental conditions.

### 4.2 Insights into the metabolism of EBPR microbial community under dynamic conditions

Our results show that the adenylate energy charge (AEC) changes dynamically throughout the EBPR cycle and may drive rapid acetate uptake and polyphosphate degradation. These findings support the hypothesis, backed by *in vitro* studies, that polyphosphate degradation is triggered by imbalances in the adenylate pool (ATP, ADP and AMP)^27,65^. Interestingly, the anaerobic phase begins with a very low AEC (Figure 2A), suggesting the cells are primed for substrate uptake and energy mobilization. The magnitude of AEC fluctuation is remarkable—previous studies on *E. coli*^*14*^ and *Synechococcus elongatus*^*66*^ report minimal changes in energy charge under variating environmental conditions. Although transient drops in AEC have been observed in *E. coli* under sudden starvation^67^, cells adapted to feast–famine regimes maintain a stable AEC^14,68^. In contrast, our data indicate that even in an adapted PAO community, AEC remains highly responsive to environmental transitions.

Notably, the most pronounced changes in AEC occurred during anaerobic substrate uptake and toward the end of the aerobic phase, while the transition between anaerobic and aerobic conditions had little immediate effect. The uridylate energy charge (UEC) showed a gradual increase under aerobic conditions, which may reflect increased biosynthetic activity, particularly in protein translation^69^. This aligns with UTP’s role in anabolic processes and supports previous reports of oxygen-induced biosynthetic activation^70^. Remarkedly, despite the major metabolic shift that occurs upon oxygen introduction—reversing PHA synthesis, and glycogen and polyphosphate degradation—the energy nucleotide pools remained relatively stable. This suggests that the switch is regulated by factors outside the adenylate and uridylate pools, possibly involving redox cofactors such as NAD(P)H/NAD(P)^+^. While these metabolites are likely central to this transition, their known instability^71,72^ poses significant challenges for accurate quantification. These findings highlight the importance of quantitative metabolomics in capturing rapid microbial responses and provide a foundation for further studies into the regulation of energy and redox metabolism in complex biofilm systems.

## 5 Conclusion

This study established and systematically evaluated a workflow for quantifying nucleotide pools in a granular biofilm microbial community performing EBPR. The stable isotope dilution mass spectrometry pipeline integrated a rapid quenching using liquid nitrogen, a boiling water extraction, and analysis by a porous graphitic carbon chromatographic separation and high resolution mass spectrometry. Application of the workflow to an EBPR anaerobic-aerobic cycle revealed dynamic changes in the adenylate energy charge (AEC) and uridylate energy charge (UEC) during acetate uptake and polyphosphate degradation, emphasizing the importance of metabolite-level regulation. In addition, the method provides a foundation to study metabolites in complex microbial biofilms.

