## Supplementary Information for "Measuring Temporal Variations of Nucleotide Pools in Microbial Granular Biofilm Performing Enhanced Biological Phosphorous Removal"

#### 1. Evaluating metabolite stability during sample analysis

The sample stability in terms of degradation was determined using the adenylate energy charge (AEC) and the nucleotide ratios of the same $^{13}C$-labeled yeast extract as proxy. For this, the $^{13}C$-labeled yeast extract was injected throughout the sample analysis at regular time intervals (Figure S1.1, S1.2, and S1.3).

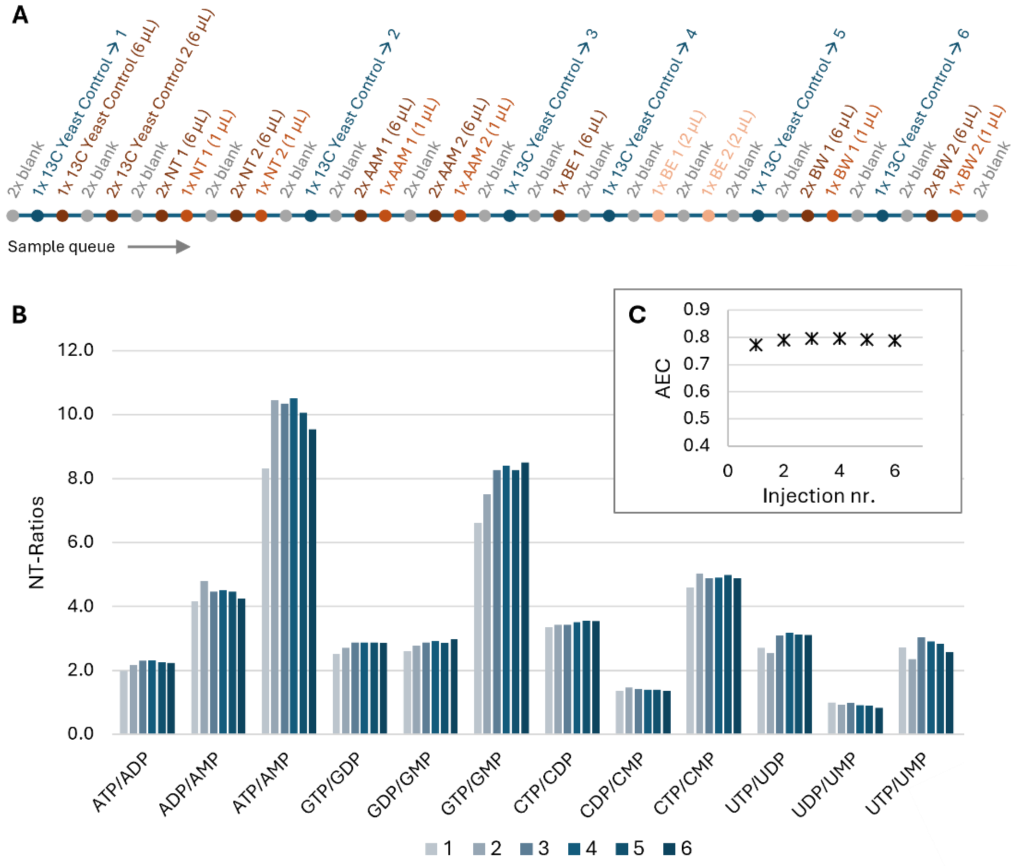

***Figure S1.1.*** **A)** PGC-HRMS sample queue. Nucleotides from a granular biofilm enrichment performing Enhanced Biological Phosphorous Removal (EBPR) were extracted employing different extraction methods: NT: no treatment, AAM: acidic acetonitrile-methanol extraction, BE: boiling ethanol extraction, BW: boiling water extraction. **B)** Nucleotide ratios of the $^{13}C$ yeast extract controls. **C)** AEC (adenylate energy charge) of the $^{13}C$ yeast extract controls. AEC = ([ATP] + 0.5*[ADP])/([ATP] + [ADP] + [AMP]). The energy charge and nucleotide ratios are calculated based on the $^{13}C$ signals of the metabolites. 1-6 corresponds to the $^{13}C$ yeast extract control injections as indicated in **A)**.

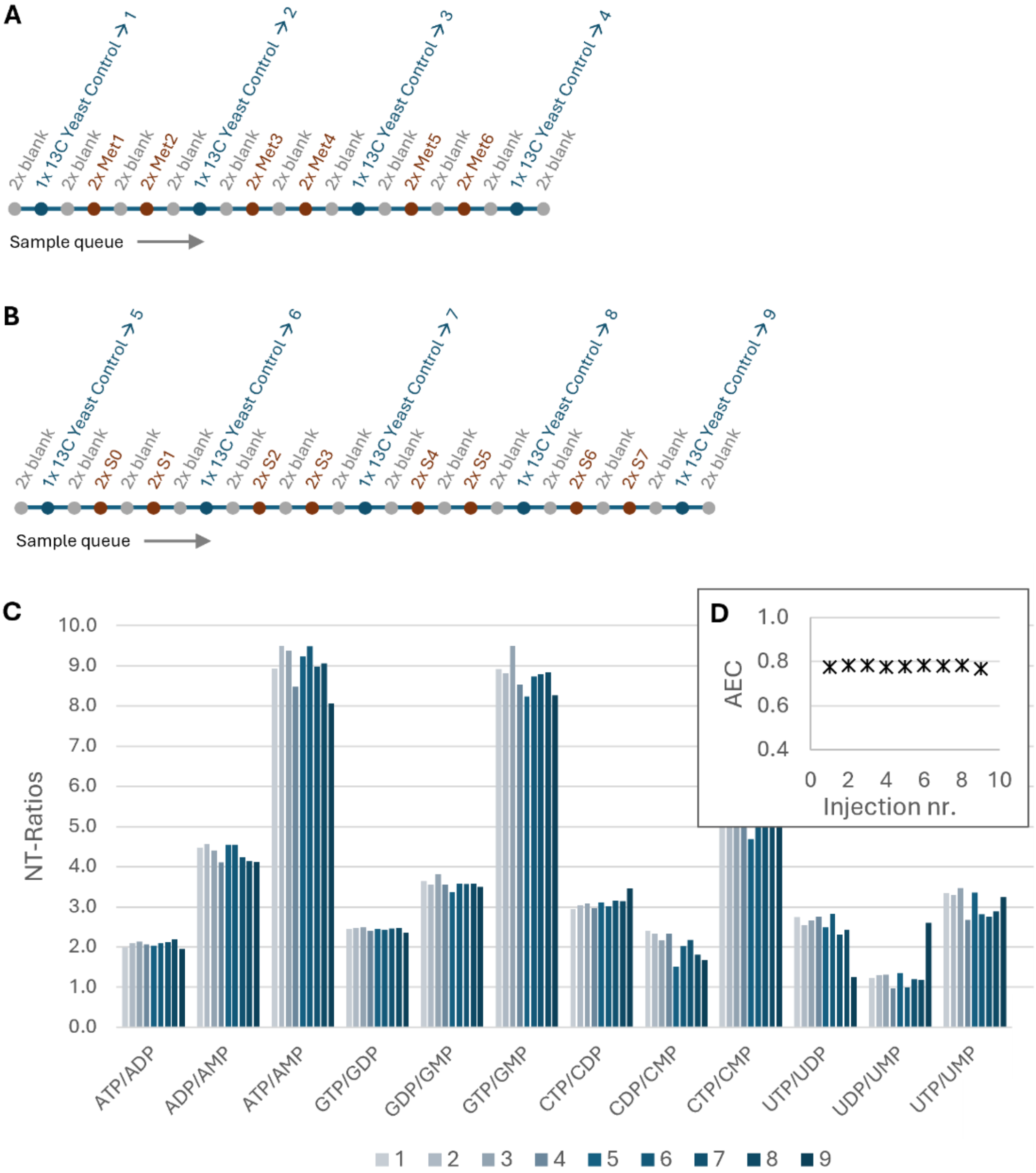

***Figure S1.2***. **A)** PGC-HRMS sample queue. Met1, Met2: Biomass from the lab-scale EBPR system was crushed in liquid nitrogen after which $^{13}$C yeast extract was added. Metabolites were extracted with the boiling water method. Met3, Met4: Biomass from the lab-scale EBPR system was crushed in liquid nitrogen, $^{13}$C yeast extract was added <u>before</u> the crushing. Metabolites were extracted with the boiling water method. Met5, Met6: Biomass from the lab-scale EBPR system was filtered using a 0.42 µm filter. The filtrate was subsequently further processed, representing the extracellular space. **B)** PGC-HRMS sample queue. S0-S7: calibration samples, mixtures of $^{12}$C metabolites of different concentrations spiked with 25 µL $^{13}$C yeast extract. **C)** Nucleotide ratios of the $^{13}$C yeast extract controls. **D)** AEC (adenylate energy charge) of the $^{13}$C yeast extract controls. AEC = ([ATP] + 0.5*[ADP])/([ATP] + [ADP] + [AMP]). The energy charge and nucleotide ratios are calculated based on the $^{13}$C signals of the metabolites. 1-9 corresponds to the injected $^{13}$C yeast extract control as indicated in **A)** and **B)**.

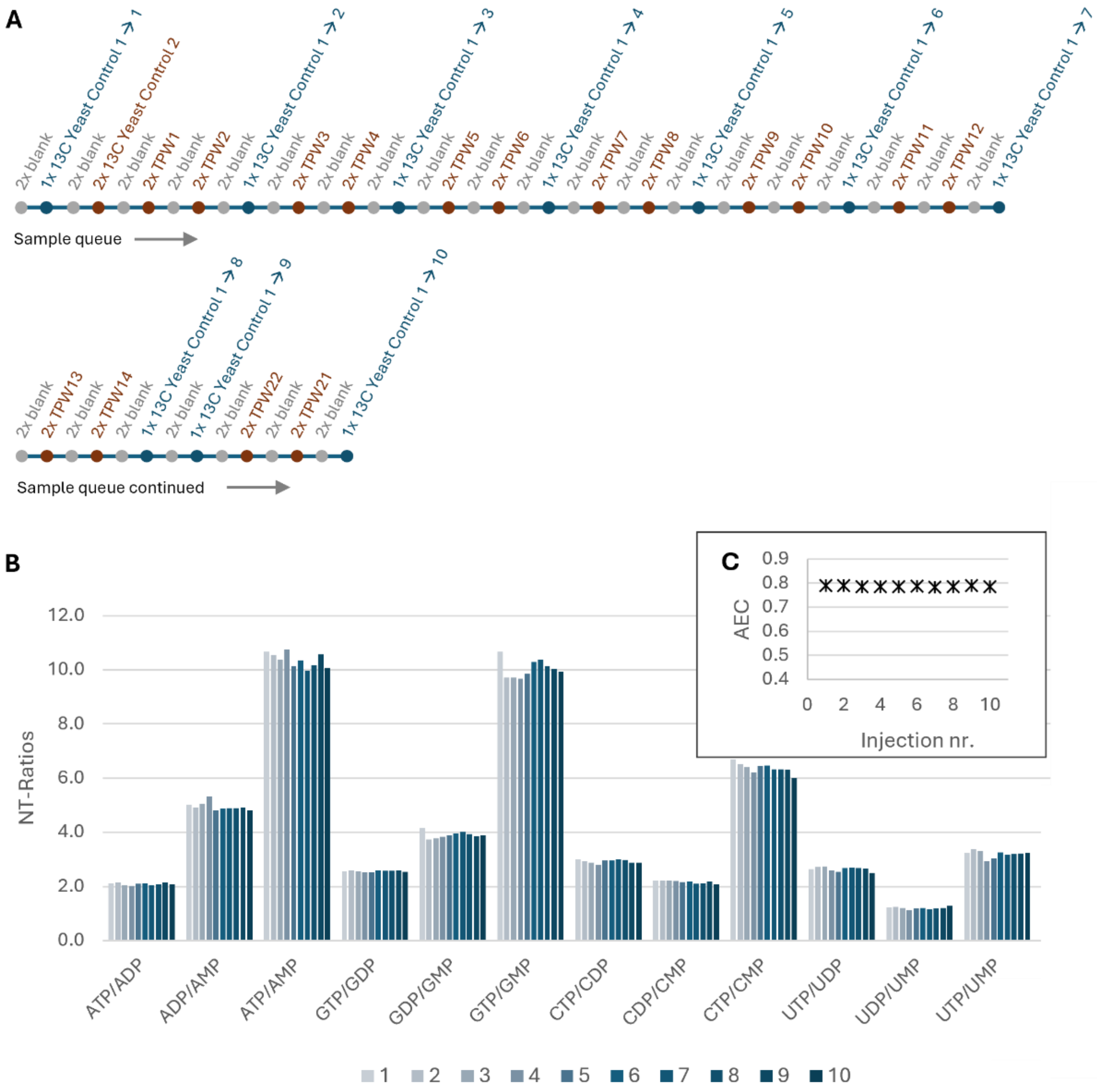

***Figure S1.3***. **A)** PGC-HRMS sample queue. TPW1-TPW14, TPW21-TPW22: metabolite extracts of the granular biofilm microbial community performing Enhanced Biological Phosphorous Removal (EBPR). **B)** Nucleotide ratios of the $^{13}C$ yeast extract controls. **C)** AEC (adenylate energy charge) of the $^{13}C$ yeast extract controls. AEC = ([ATP] + 0.5*[ADP])/([ATP] + [ADP] + [AMP]). The energy charge and nucleotide ratios are calculated based on the $^{13}C$ signals of the metabolites. 1-10 correspond to the injected $^{13}C$ yeast extract control as indicated in **A)**.

#### 2. Nucleotide ratios of spiked 13C yeast after employing different extraction protocols

The nucleotide ratios of the $^{13}$C-labeled yeast extract that was spiked to the granules from the lab-scale EBPR system, and which were extracted with different extraction protocols, were compared to the $^{13}$C-labeled yeast extract control (Figure S2).

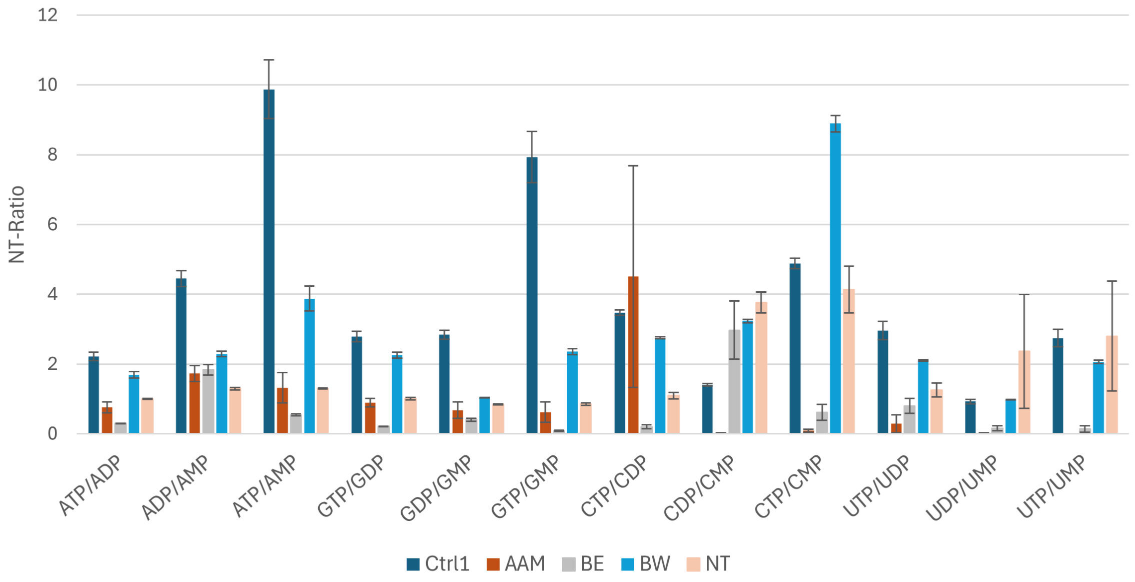

***Figure S2***. Nucleotide ratios of the spiked $^{13}$C yeast extract after employing different extraction protocols to granules from the lab-scale EBPR system. AAM: acidic acetonitrile-methanol extraction, BE: boiling ethanol extraction, BW: boiling water extraction, NT: no treatment. Ctrl1: untreated $^{13}$C yeast extract.

#### 3. Nucleotide areas of extracted granules from lab-scale EBPR system

The nucleotide signals of the extracted granules from the lab-scale EBPR system were compared to determine the extraction efficiency of different extraction protocols (Figure S3). No significant amount of $^{12}C$ metabolites were observed in the uniformly $^{13}C$-labeled yeast extract control, which confirms, that all extracted metabolite signals originated from the EBPR microbes. Interestingly, only trace signals were observed for CTP, CDP, CMP, UTP, UDP, and UMP in the samples extracted with the acidic acetonitrile-methanol (AAM) method and the boiling ethanol (BE) method.

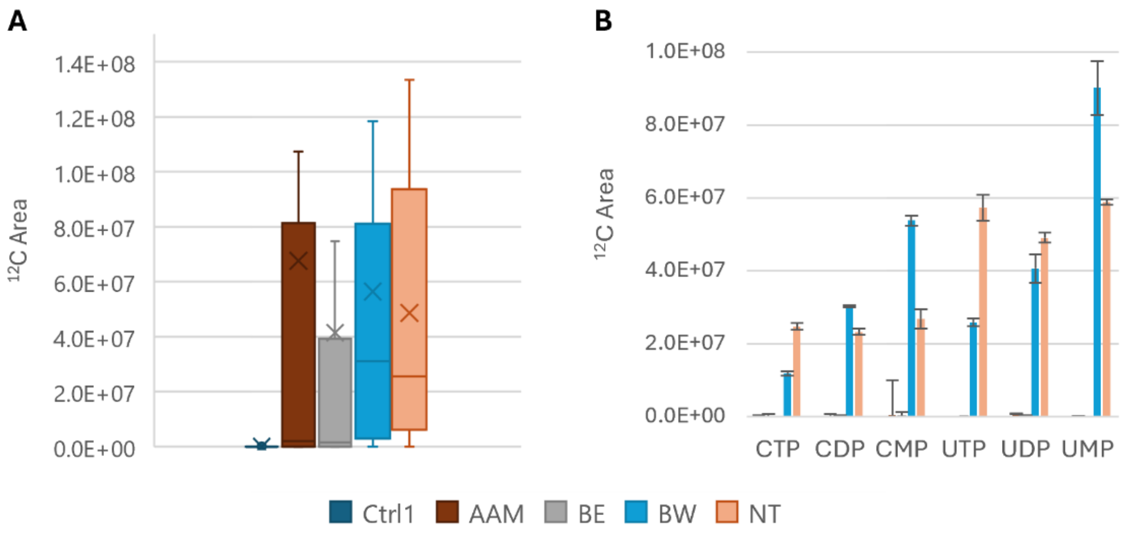

***Figure S3***. Differently extracted granules from the lab-scale EBPR system. AAM: acidic acetonitrile-methanol extraction, BE: boiling ethanol extraction, BW: boiling water extraction, NT: no treatment. Ctrl1: uniformly $^{13}C$-labeled yeast extract. **A)** Boxplot of the summed $^{12}C$ nucleotides. **B)** Uridine and cytidine nucleotide signal intensities.

#### 4. Nucleotide calibration curves

Calibration curves were established for all nucleotides by plotting the $^{12}C/^{13}C$ HRMS signal against the known $^{12}C$ nucleotide concentrations (Figure S4) to allow absolute quantification of the extracted nucleotides from the microbes in the EBPR system.

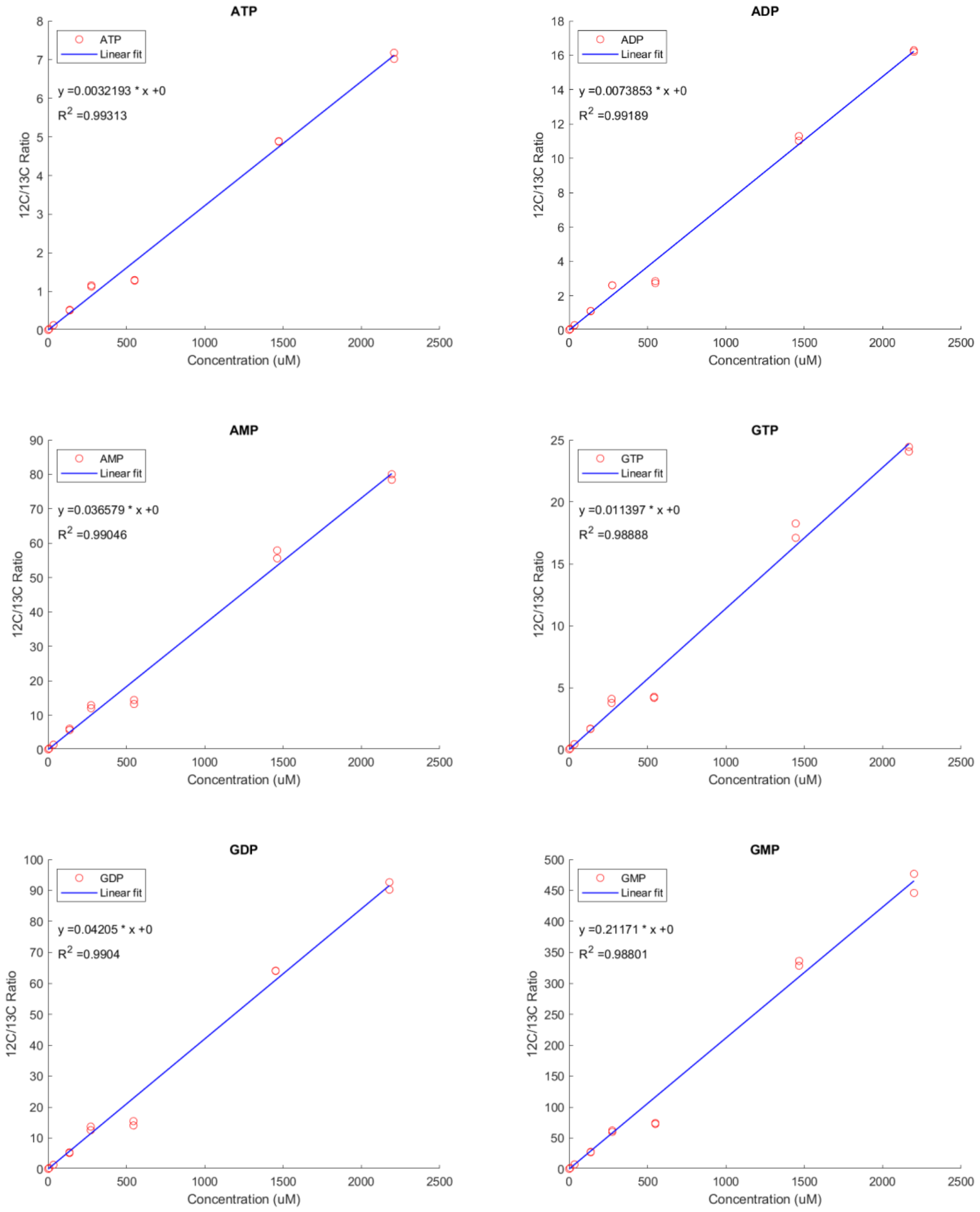

***Figure S4***. Calibration curves of the nucleotides AxP, GxP, CxP and UxP. The $^{12}C/^{13}C$ HRMS signal is plotted against the known $^{12}C$ nucleotide concentrations.

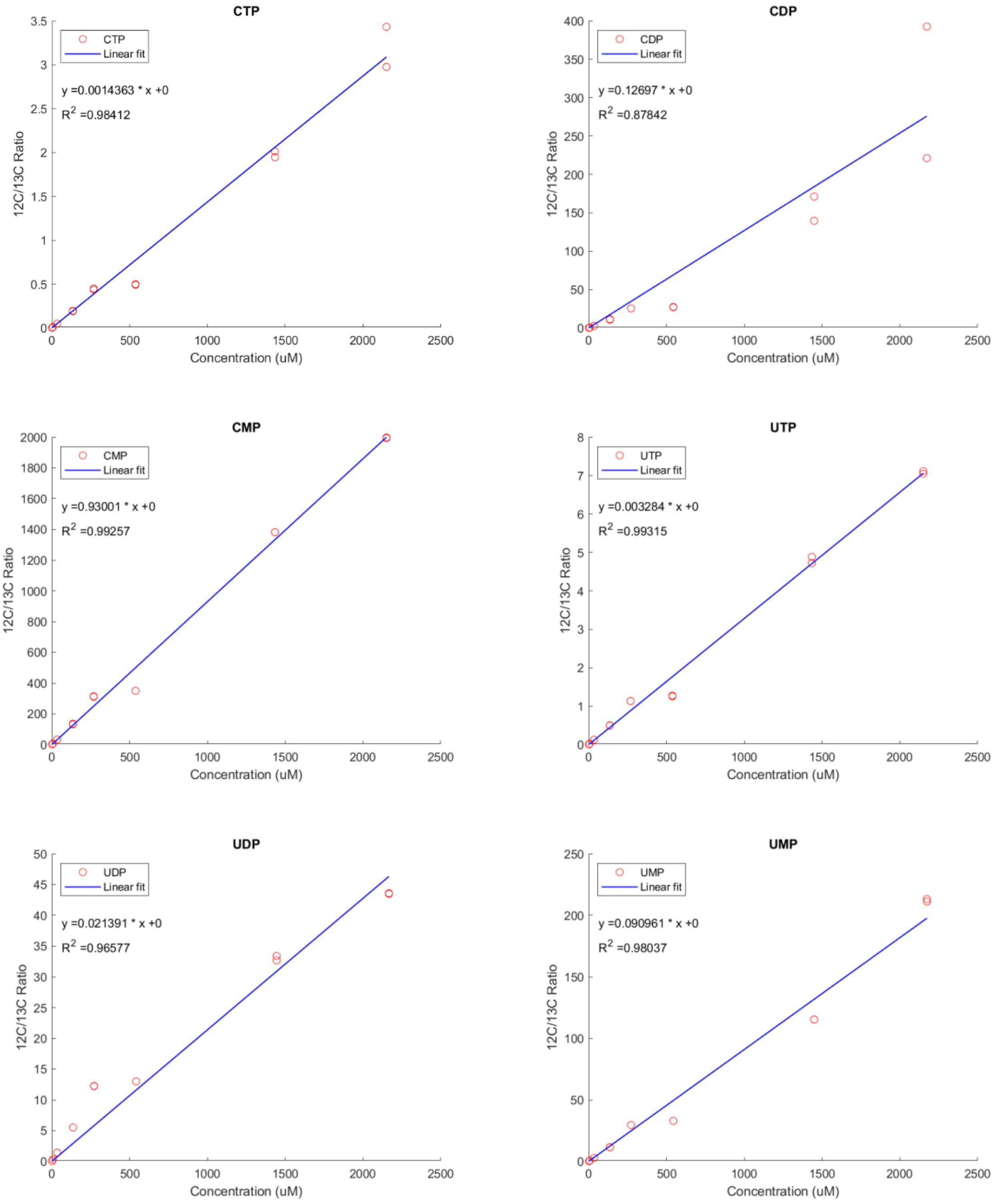

***Figure S4 continued***. Calibration curves of the nucleotides AxP, GxP, CxP and UxP. The $^{12}C/^{13}C$ HRMS signal is plotted against the known $^{12}C$ nucleotide concentrations.

### 5. Nucleotide elution profiles on porous graphitic carbon (PGC)

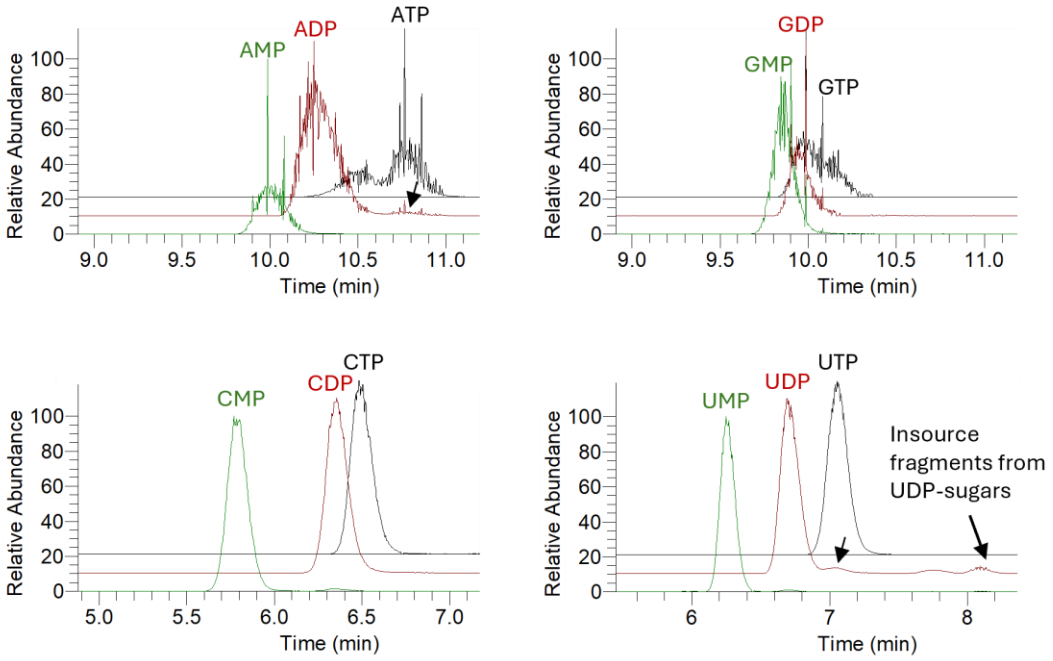

***Figure S5.1***. Extracted Ion chromatograms (XIC) of the $^{12}C$ nucleotides from a metabolite extract of a granular biofilm microbial community performing enhanced biological phosphorus removal (EBPR ). Analysis was performed by high resolution mass spectrometry in negative mode, using porous graphitic carbon (PGC) as separation phase. The masses were extracted with a 5 ppm error range. Arrows indicate insource fragments.

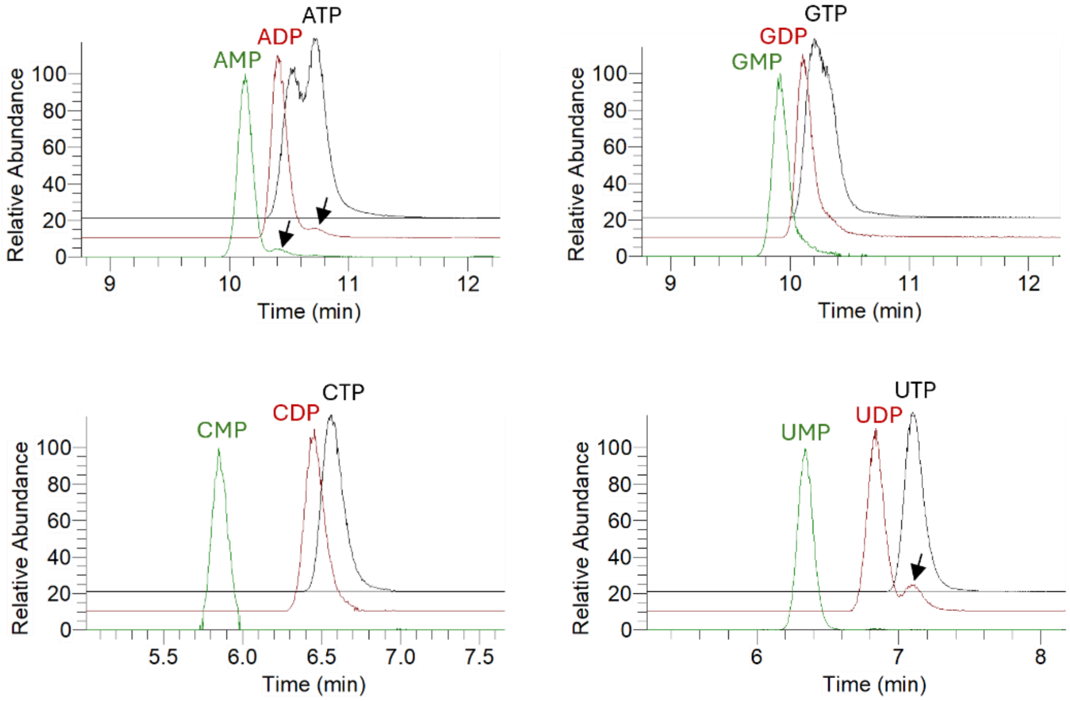

***Figure S5.2***. Extracted Ion chromatograms (XIC) of the $^{13}C$-labelled nucleotides in a uniformly $^{13}C$-labelled yeast extract measured by high resolution mass spectrometry in negative mode, using porous graphitic carbon (PGC) as separation phase. The masses were extracted with a 5 ppm error range. Arrows indicate insource fragments.

#### 6. Metagenome taxonomic profiling

Whole metagenome sequencing was performed followed by taxonomic classification of raw sequencing reads using Kraken2 (Figure S6).

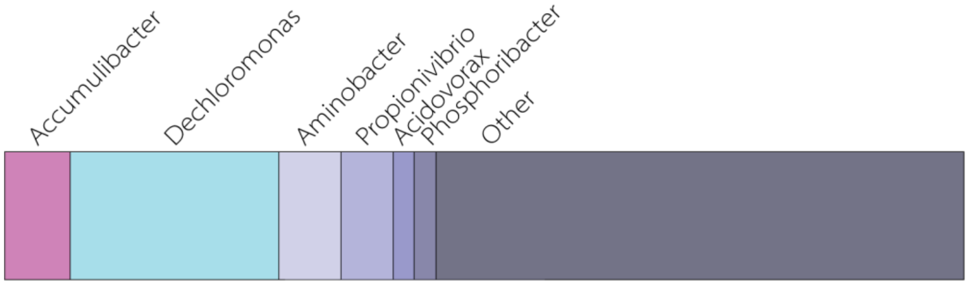

***Figure S6***. Microbial composition of the granular biofilm performing enhanced biological phosphorus removal (EBPR) based on taxonomic classification of raw sequencing reads obtained from whole metagenome sequencing.
